# Parvalbumin-expressing interneurons improve sensory discrimination by shaping noise geometry in primate V1

**DOI:** 10.64898/2026.06.11.731577

**Authors:** Sandra Cole, David G.C. Hildebrand, Lauri Nurminen

## Abstract

The ability to discriminate sensory stimuli is fundamental to sensory-guided behavior. Sensory discrimination by neural populations is constrained by the magnitude of trial-to-trial spiking variability, or noise, and its geometric alignment with the stimulus encoding directions of the population response. Computational models predict that cortical inhibition shapes the noise geometry, but causal evidence from primate cortex is lacking. We used optogenetic stimulation of parvalbumin-expressing inhibitory interneurons (PV-cells) combined with high-density extracellular recordings in primary visual cortex of awake marmoset monkeys to test this prediction. Stimulating PV-cells improved the discriminability of V1 population responses, without expanding the stimulus-related signal magnitude. Instead, PV-cell activation compressed shared trial-to-trial variability and rotated it away from the stimulus-coding direction. Our results establish a causal role for inhibition in shaping the geometry of neural population responses to improve sensory discrimination in the primate visual cortex.

---

The ability to discriminate between sensory stimuli is fundamental to perception and cognition, and sensory scientists have searched for the principles underlying this ability for more than a century^1^. Since the earliest psychophysical observations, formalized in signal detection theory^2^, it has been recognized that discrimination performance is bounded by stochastic fluctuations in the internal variables that represent sensory stimuli. These internal variables are carried by the spiking activity of neural populations, which is variable between presentations of the same stimulus^3,4^, and can thus limit behavioral performance^5–8^. Neural discrimination performance is constrained by two geometric aspects of the variable population spiking activity. Signal geometry describes how distinct stimuli are mapped onto population response patterns^9^, while noise geometry describes the structure of the shared, trial-to-trial spiking variability across the population^10^. Theoretical studies predict that the relative alignment of signal and noise determines how much the noise matters for discriminability^11^.

When noise is aligned with the signal, as suggested by previous studies^12^, a downstream decoder cannot distinguish stimulus-driven responses from noise fluctuations, and discrimination suffers. When the noise is orthogonal to the signal, even substantial variability may not affect discrimination. Improving sensory discrimination therefore requires geometric reshaping of the noise relative to the signal.

Cognitive processes such as attention^13–15^, memory^16^, and learning^17^ are known to improve behavioral discrimination performance and reduce shared spiking variability in sensory cortex. While these behavioral and neural effects are well established, the circuit mechanisms that implement them remain largely unknown. Converging evidence from computational studies, spanning abstract rate models^18^, balanced networks^19^, and large-scale spiking networks with tens of thousands of neurons and realistic spatial connectivity^20^, points to inhibition as the culprit, with several models proposing that cognitive processes act by recruiting inhibitory interneurons^21^. Consistent with this view, experimental studies have reported correlations between the activity of putative interneurons, variability, and behavioral performance^15^. However, this evidence is correlative, and causal tests of how inhibition affects spiking variability have been missing.

Here, we establish a causal role for cortical inhibition in controlling the shared variability of neural population responses in the primate visual cortex, and show that inhibition improves discrimination through geometric reorganization of spiking variability. We combined optogenetic stimulation of parvalbumin-expressing inhibitory interneurons (PV-cells) in primary visual cortex (V1) of awake, fixating marmoset monkeys with high-density extracellular electrophysiological recordings. This enabled us to study neural population spiking activity while manipulating the responses of PV-cells. Using covariance-regularization techniques to account for the limited number of trials in primate electrophysiology, we find that stimulating PV-cells compresses the shared component of neural noise and reduces its alignment with the stimulus-related signals. Together, these changes enhance the discriminability of stimuli in primate V1. Our results identify PV-cells as a circuit-level mechanism for shaping population spiking activity, and provide a causal counterpart to the long-standing correlative link between interneuron activity, shared spiking variability, and discrimination performance.

## RESULTS

We used optogenetic stimulation of parvalbumin-expressing inhibitory interneurons (PV-cells) combined with high-density extracellular electrophysiological recordings in two awake marmoset monkeys to learn how inhibition affects coding of visual information in primate V1.

### Optogenetic stimulation of PV-cells in awake marmosets

The animals were first trained to maintain stable fixation on a small (0.6 – 0.8°) centrally presented marmoset face in exchange for a juice reward. This establishes the behavioral control required for subsequent visual stimulation experiments. Once reliable fixation performance was achieved, a recording chamber was surgically implanted over the primary visual cortex to provide repeated access for electrophysiology and optical stimulation. Through small openings made in the dura mater visualized under a microscope within the chamber, viral vectors (AAV.PhPeB.S5E2.ChR2-mCherry^38^) carrying the genes for the excitatory opsin Channelrhodopsin were injected to drive selective expression in parvalbumin-expressing (PV+) interneurons. After allowing at least 4 weeks for viral expression, neural responses to natural images sampled from the Berkeley segmentation dataset were recorded with Neuropixels NHP 1.0 multielectrode arrays simultaneously with optical stimulation of PV-cells.

**Figure 1.**
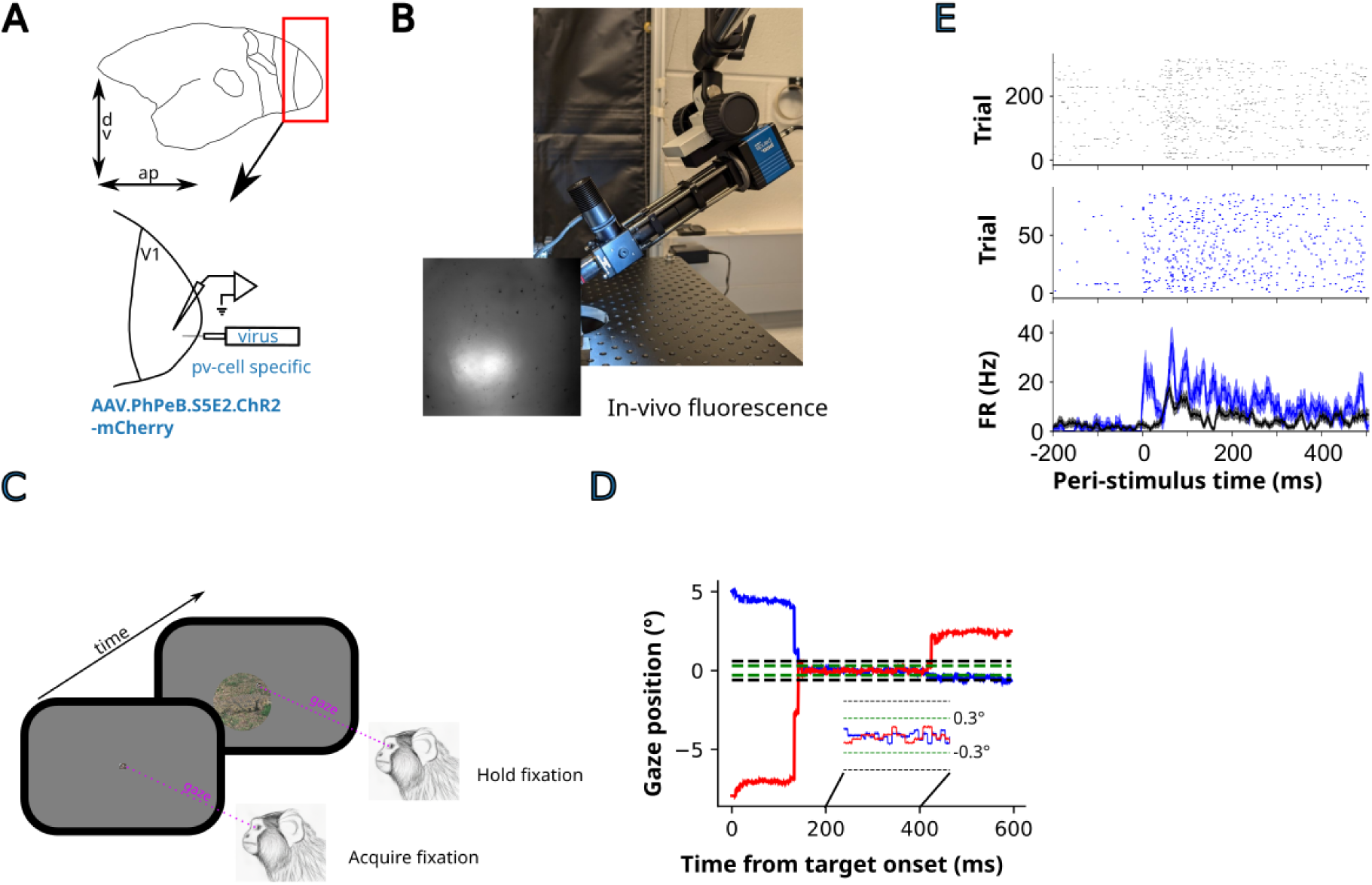
Optogenetic stimulation of PV-cells in awake marmosets: **A)** Schematics of virus vector injections and electrophysiological recordings in marmoset V1. **B)** In-vivo fluorescence was used to target and time Neuropixels multielectrode array recordings. **C)** Illustration of the awake, fixating paradigm and the stimuli used. **D)** Gaze traces in an example trial. **E)** An example cell directly driven by optogenetic stimulation.

### PV-cells improve visual stimulus discrimination

To understand how PV-cells affect the coding of visual information in primate cortex, we asked whether activating them changes the ability of neural populations in V1 to discriminate stimuli. Because population responses are high-dimensional, we first used partial least squares (PLS) dimensionality reduction to identify the dimensions of neural population response space that best separated the stimuli^12^. Projected onto the first two PLS dimensions, the stimulus-evoked responses were more separated when PV-cells were stimulated than in the control condition (Figure 2A, B). The same effect was apparent along Fisher’s linear discriminant, the one-dimensional axis for the largest stimulus separation, where PV-cell stimulation pushed the two response distributions apart (Figure 2C, D). These analyses suggest that PV-cell activation improves discrimination in primate V1, which motivates a quantitative analysis across sessions.

**Figure 2.**
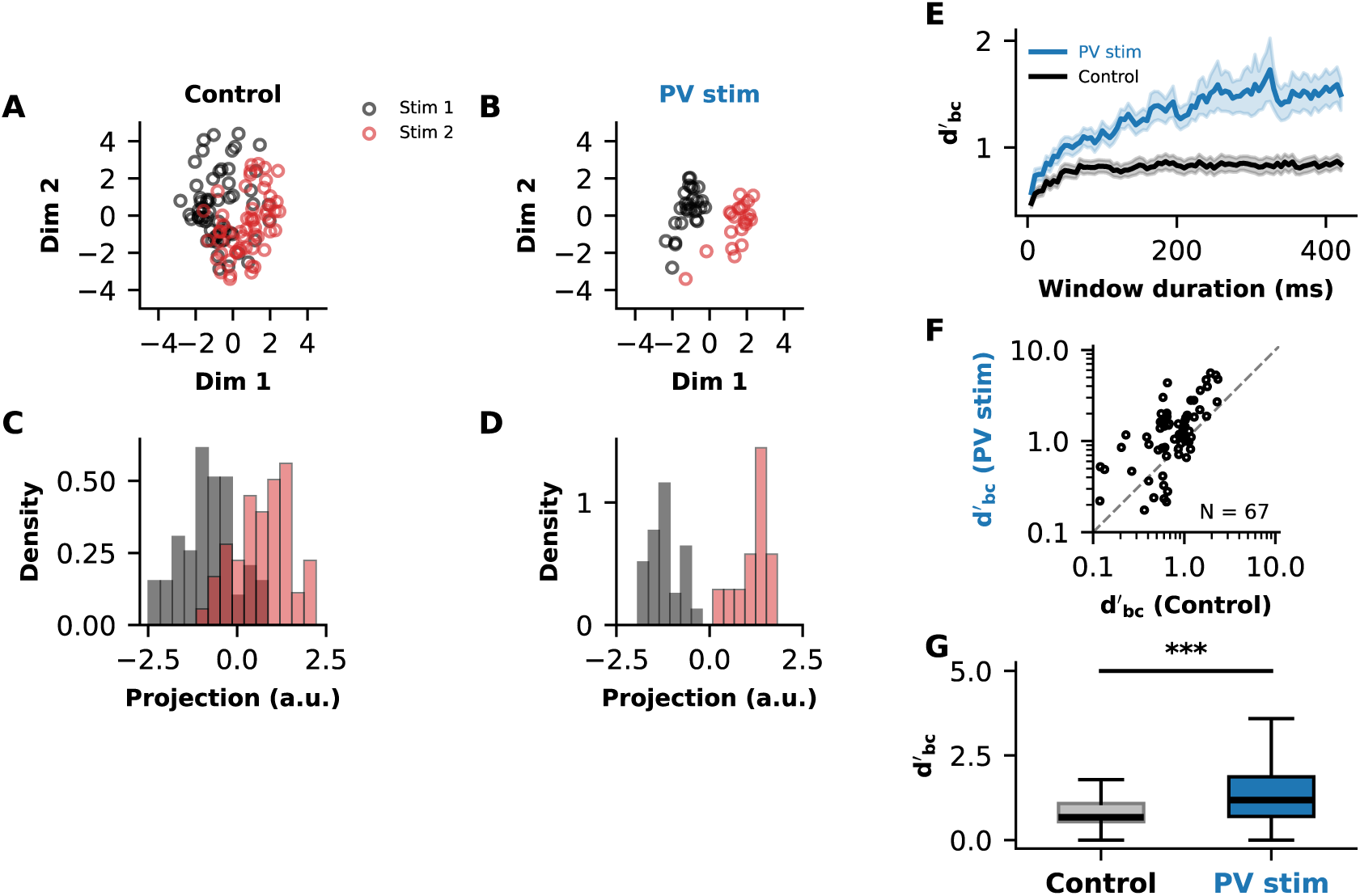
PV-cells improve stimulus discrimination: **A)** Neural population response to two natural stimuli projected onto the first two PLS dimensions. **B)** The same as A except for PV-cell stimulation trials. **C)** The population response in A projected onto Fisher’s linear discriminant axis. **D)** The same as C except for stimulation trials. **E)** Bias-corrected discriminability as a function of the spike-count window duration, averaged over sessions. **F)** Bias-corrected discriminability across all sessions and stimulus pairs. **G)** Distributions of bias-corrected discriminability. The black horizontal lines mark the medians of the distributions. *** = bootstrap-test for the difference between the medians p < 0.001.

To quantify how PV-cell stimulation affects neural coding across all sessions and animals, we computed the bias corrected discriminability index (d′_bc_) of an optimal linear decoder applied to the population responses^22^. Because changes in firing rate can trivially inflate d′ and confound comparisons across conditions, we matched the mean spike-count distributions between trials with and without PV-cell stimulation, and corrected for differences in trial counts between conditions. We found that d′_bc_ grew with the duration of the spike-count window, as expected from temporal accumulation of stimulus evidence, and at longer integration windows PV-cell stimulation yielded significantly higher mean d′_bc_ than the control condition (Figure 2E; mean ± s.e., control 0.84 ± 0.07 vs PV stim 1.53 ± 0.16, paired t-test p=2.5×10^-7^]). This enhancement was consistent across the majority of experimental sessions and stimulus pairs (Figure 2F) and produced a significantly higher median d′_bc_ during PV-cell stimulation at the population level (Figure 2G; control median = 0.67 [95% CI 0.60, 0.92], PV stim median = 1.18 [95% CI 0.94, 1.54, paired bootstrap test p=9.99×10^-5^]). Together, these results show that activation of PV-cells improves the capacity of primate V1 neural populations to discriminate visual stimuli. In the remainder of the study, we will investigate how PV-cells shape neural population spiking to improve discriminability.

### Signal magnitude does not underlie improved discriminability

PV-cell stimulation could improve discriminability either by pushing the mean responses to different stimuli further apart, i.e. by increasing signal magnitude, by changing the noise, or a combination of the two. We asked whether the increased discriminability could be accounted for by an increase of the signal magnitude. For each session, we computed the Euclidean distance between the mean population-response vectors of every stimulus pair in the control condition and when PV-cells were stimulated. Across sessions and animals, PV-cell stimulation did not produce a systematic change in the signal magnitude, for some stimuli and sessions the signal mangnitude increased, for others it decreased, and there was no statistically signifcant effect at the population level (Figure 3A-C, pair level tests were performed with two-sided Wilcoxon signed rank test p = 0.76, per penetration tests again ΔD = 0 with one-sample Wilcoxon signed rank test p = 0.68). Thus, we next ask if PV-cells increase stimulus discriminability via spiking variability.

**Figure 3.**
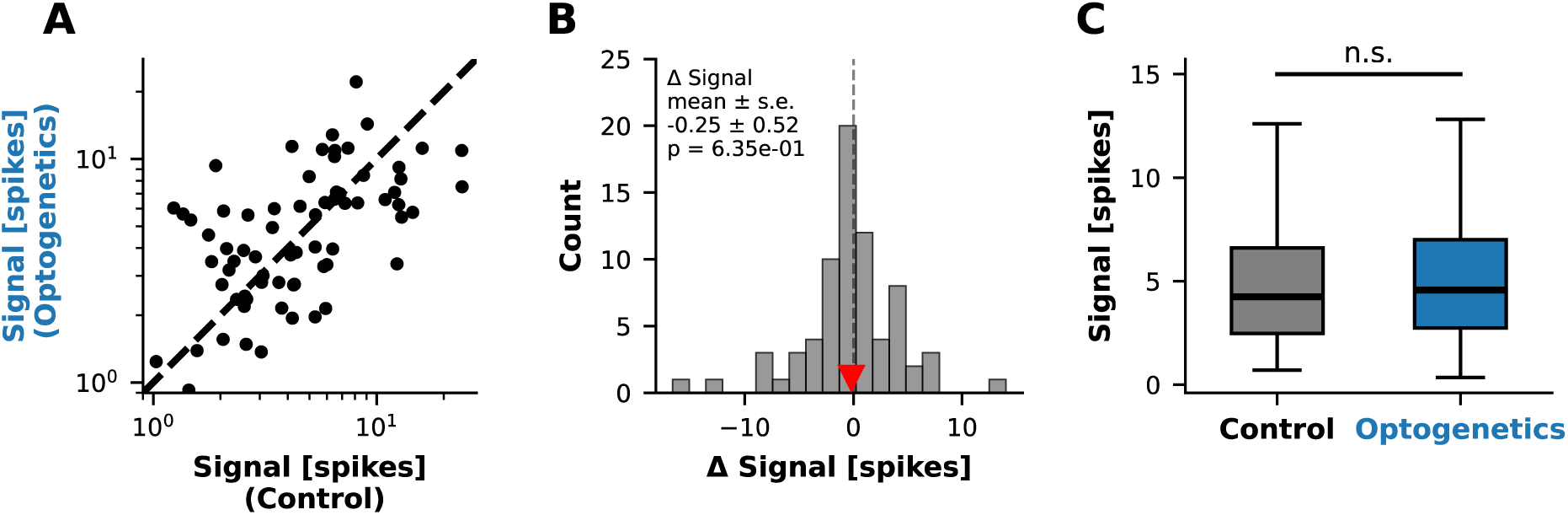
Signal magnitude does not underlie improved discriminability: **A)** Signal magnitude. **B)** Change in the signal magnitude with PV-cell stimulation. The p-value refers to t-test for Δ Signal = 0. **C)** Distribution of the signal magnitude with and without PV-cell stimulation. n.s. refers to the non-significance of bootstrap test for the difference between the medians.

### PV-cells reduce single-cell spiking variability

To understand how PV-cell stimulation affects spiking variability in V1, we analyzed Fano-Factor (FF), the ratio of spike-count variance to mean across repetitions of the same stimulus. Fano-Factor was computed in a sliding 50 ms window, separately for control and stimulation trials. Stimulating PV-cells reduced Fano-Factor throughout the response, but the reduction was largest shortly after the onset and attenuated over the course of the response (Figure 4A), and the reduction was consistent across the population and PV-cell stimulation decreased variability in the majority of the cells (Figure 4B). Quantifying variability in a 400 ms count window (100–500 ms from the stimulus onset), matched to the count window duration at which d′_bc_ was maximally increased (Figure 2), showed that PV-cell stimulation significantly reduced the median Fano-Factor across the population (Figure 4C; median ± 95% CI, 1.48 ± [1.40, 1.55] during control vs. 1.29 ± [1.24, 1.34] during PV stim). Thus, recruiting PV-cells reduces trial-to-trial variability so that stimulus driven responses tightly fluctuate around their mean.

**Figure 4.**
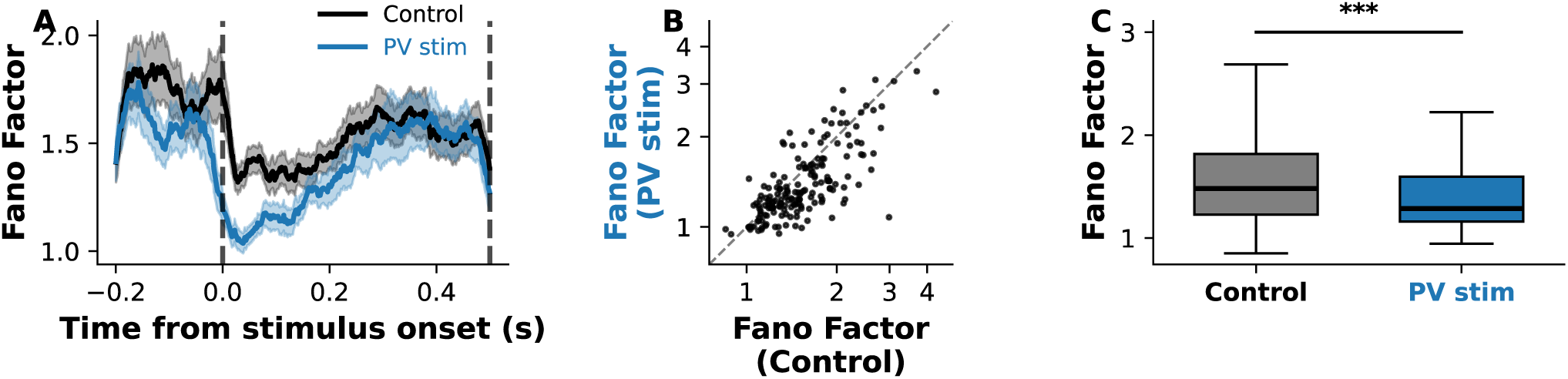
PV-cells reduce single-cell spiking variability: **A)** Temporal evolution of fano-factor computed in a 25ms sliding window. **B)** Fano-factor of all single-units recorded for this study. **C)** Distributions of the fano-factor. *** = bootstrap-test for the difference between the medians p < 0.001.

The Fano-Factor analysis is consistent with the idea of PV-cells improving discriminability by compressing noise around each stimulus representation. However, Fano-Factor is a single-neuron measure that conflates private variability that is independent across neurons, and shared variability that is correlated across the population^23^. This distinction matters because private, but not shared, variability can be averaged out. Thus, to better understand how PV-cells improve discriminability of visual information in primate cortex, we next asked how stimulating PV-cells affects shared variability in V1.

### PV-cells decorrelate spiking variability

To understand how PV-cells affect shared spiking variability in V1 neurons, we estimated pairwise noise correlations between simultaneously recorded neurons. Estimating large correlation matrices from the limited number of trials available in awake primate recordings is an ill-posed problem. When there are more neuron pairs than independent observations, sample covariance matrices are unstable and biased, and individual correlation estimates become noisy. To overcome this problem, we used an approach from previous studies^24–26^ and regularized each covariance matrix by decomposing it into a low-dimensional shared component and an independent per-neuron component using factor analysis (FA). The number of factors was selected by cross-validation and all correlations below are derived from these regularized covariance matrices. To give an unfiltered view of the effect of PV-cell stimulation on shared trial-to-trial variability, the two example neuron pairs shown for illustration (Figure 5A, B) were computed directly from the z-scored spike counts.

**Figure 5.**
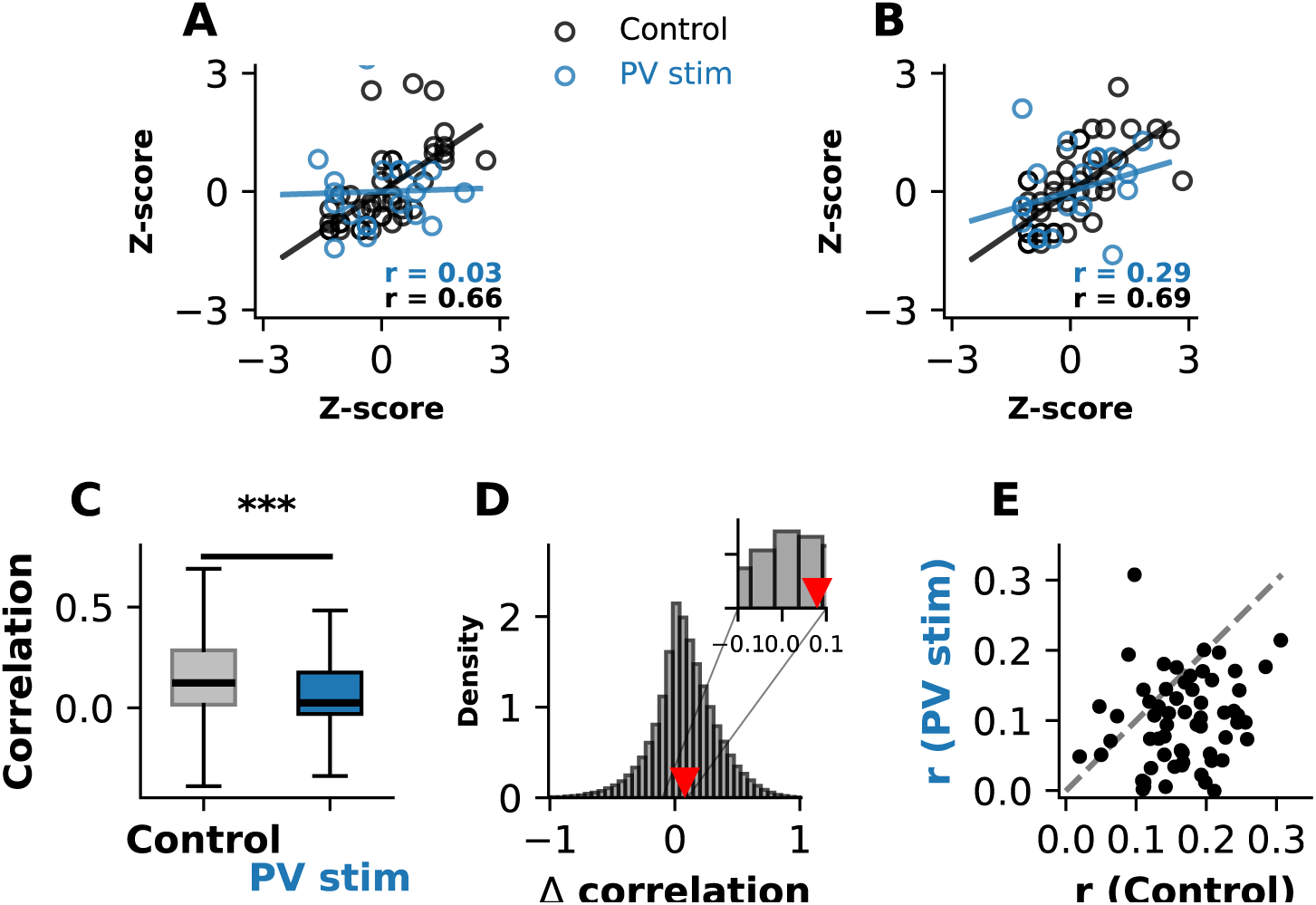
PV-cells decorrelate spiking variability: **A, B)** Z-scored responses of two unit pairs. **C)** Distributions of noise correlations. *** = bootstrap-test for the difference between the medians p < 0.001. **D)** Change in noise correlations with PV-cell stimulation. Here positive values indicate that correlations were higher in the control condition. **E)** Noise correlations averaged over sessions.

Figures 5A and B shows the z-scored single-trial responses of two example neuron pairs to repeated presentations of the same stimulus. In the control condition (black), the responses of both pairs were correlated from trial to trial: when one neuron of a pair fired above its mean rate, the other tended to do the same. Stimulating PV-cells reduced the correlations across trials. These examples suggest that PV-cell recruitment decorrelates spiking variability in V1.

To quantify this effect across the entire dataset, we computed pairwise spike-count noise correlations from FA-regularized covariance matrices for every session, stimulus, and recorded cell pair. The distribution of pairwise correlations was significantly shifted toward zero when PV-cells were stimulated, compared with the control condition (Figure 5C; median ± 95% bootstrapped CI, control 0.1237 ± [0.1230, 0.1242] vs. PV-cell stimulation 0.0253 ± [0.0250, 0.0255] bootstrap test for the difference between the medians, p < 0.0001). On average, stimulating PV-cells reduced pairwise correlations by ∼0.07 (Figure 5D). This is a substantial change corresponding to a roughly 50% reduction in average correlation between pairs. To confirm that the effect was seen across sessions, we averaged the correlations within each penetration and stimulus and compared the resulting averages with and without PV-cell stimulation (Figure 5E). This analysis showed that PV-cell stimulation consistently reduced the average correlated spiking variability across sessions and stimuli (mean ± s.e. across sessions and stimuli, control vs PV-cell stimulation, 0.16 ± 0.007 vs 0.10 ± 0.008, independent samples t-test p=1.03×10^-8^). Thus, our data shows that PV-cell stimulation decorrelates spiking variability in primate V1.

These analyses establish that stimulating PV-cells reduces the magnitude of shared variability in V1, and thus extends the body of correlative work linking inhibition, noise correlations, and discrimination performance. However, these analyses do not reveal whether PV-cells improve discriminability through changes in the private or the shared noise.

### Noise components underlying PV-enhanced discrimination

To understand how the different noise components related to improved discrimination under PV-cell stimulation, we computed discriminability of decoders in which all the other components were estimated from the control trials, but the private or the shared noise was estimated from the PV-cell stimulation trials (Figure 6). This clarifies the contribution of each noise component to the improved discriminability.

**Figure 6.**
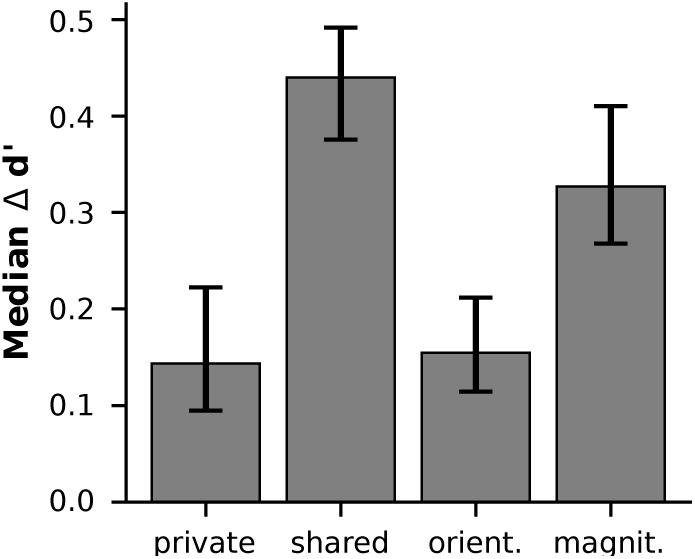
Noise components underlying PV-enhanced discrimination: Median change in d-prime, the errors show 95% bootstrapped CI, private = private noise from PV-trials, shared = shared noise from PV-trials, orient. = the eigenvectors of the noise covariance from PV-trials, magnit. = the spectrum of the noise covariance from PV-trials.

Using the private noise from the PV-cell stimulation trials increased d-prime by (median ± 95% CI) 0.14 ± [0.09, 0.22] (Wilcoxon signed-rank test on the paired differences p = 10^-4^) and using the shared noise from PV-cell stimulation improved discriminability and increased d-prime by (median ± 95% CI) 0.44 ± [0.38, 0.49] (Wilcoxon signed-rank test on the paired differences p = 10^-4^). This shows that most of the improvement in discriminability was due to the effect of PV-cells on the shared spiking variability.

PV-cells could improve discriminability either by compressing the magnitude of the shared noise or by rotating it so that it interferes less with the signal. To learn which one of these possibilities is true, we computed discriminability under decoders in which the eigenvectors of the share noise covariance were estimated from the PV stimulation trials, and all the other components, including the eigenvalues of the noise covariance, were estimated from control trials. This analysis reveals how much PV-cell induced rotation of the shared noise improves discriminability. Using eigenvectors from the PV stimulation trials increased d-prime by (median ± 95% CI) 0.15 ± [0.11, 0.25] (Wilcoxon signed-rank test on the paired differences p = 10^-4^). To understand how much change in the shared noise magnitude improved discriminability, we used a decoder in which the eigenvalues values of the noise covariance were estimated from PV stimulation trials, and all the other parameters from control trials.

Under this decoder, d-prime increased by (median ± 95% CI) 0.33 ± [0.27, 0.41] (Wilcoxon signed-rank test on the paired differences p = 10^-4^).

These analyses show that PV-cells improve discriminability mainly by compressing the shared noise, with the orientation of the shared noise and the private noise having smaller, but statistically significant, effects.

## DISCUSSION

We used optogenetic stimulation of parvalbumin-expressing inhibitory interneurons in V1 of awake marmoset monkeys, combined with high-density extracellular recordings and covariance regularization techniques, to study the neural circuit mechanisms that control stimulus discriminability in primate V1. Stimulating PV-cells improved stimulus discriminability and the effect was not explained by increased magnitude of the stimulus-related signal.

Instead, stimulating PV-cells reduced trial-to-trial variability in single neurons and reduced the magnitude of shared variability between neurons. Most of the improvement was due to the effect of PV-cells on the magnitude of the shared spiking variability, with a smaller effect due the orientation of the shared variability. Taken together, these results establish a causal role for cortical inhibition in improving stimulus discriminability through shaping the geometry of spiking variability in neural populations of the primate visual cortex.

Cognitive processes such as attention, memory and learning improve sensory performance and reduce shared neural response variability in the cortex ^13–15,17^. Computational models ranging from abstract rate-models^18^ to spiking neural networks^21^ suggest that inhibition underlies these effects. Similarly, V1 neurons’ receptive field surround affects correlated variability^26–28^, which hints toward inhibitory control. Until now, the most direct experimental evidence for the role of inhibition has been indirect and derived from correlating the activity of electrophysiologically identified fast-spiking cells (putative parvalbumin-expressing cells), with behavioral performance and shared variability^13,14^. By using this strategy, Mitchell et al. showed that spatial attention decreased trial-to-trial variability across cell-types and increased the firing-rate of putative inhibitory cells more than broad-spiking cells. Similarly in the sensory cortex of different rodent species^29^, electrophysiologically identified fast-spiking cells are more active during epochs of weakly correlated firing. This correlative evidence suggests that increased PV-cell activity dampens the variability of cortical responses. Here, by stimulating PV-cells in the marmoset visual cortex, and measuring the consequences of this manipulation on neural responses, we established a causal role for inhibition in improving discriminability and reducing shared variability of spiking activity in V1. While we did not measure behavior, the fact that PV-cell stimulation reproduced known effects of cognitive processes on neural population responses, i.e. reduced shared variability and improved discrimination performance, lends support for the hypothesis that cognitive processes affect sensory processing through inhibition.

Spiking neural network models of the visual cortex predict that increased inhibitory drive shifts the network into a more strongly stabilized regime and dampens shared spiking variability^21^.

Consistent with this prediction, PV-cell stimulation in our data quenched shared variability. At the single-cell level, PV-cells mediate gain control or normalization both in rodents^30^ and primates^31^, and recent work has reported a negative correlation between single-cell variability and the strength of normalization^32^. Together, these mechanisms provide a parsimonious account of how PV-cells dampen shared variability in primate V1.

Our data suggests that PV-cells rotate the shared noise relative to the signal, and that this rotation plays a small but significant role in improving discriminability. The rotation can be understood as a geometric consequence of activity-dependent divisive gain control by PV-cells. The activity of PV-cells is proportional to the local network activity^33^ and they provide the strongest inhibition to the pyramids with the largest stimulus-driven responses^30,34^. These are the cells that can at least in principle, depending on their tuning, contribute most to the signal in the population response space. The variance of these cells is therefore preferentially attenuated, which shrinks the shared-noise covariance more strongly along the signal direction than along directions orthogonal to it, which rotates the shared noise. Thus, the same PV-cell inhibition that controls single-cell gain^30,31^ underlies, at the population level, compression and the rotation of spiking variability.

Recent studies have aimed at answering whether sensory information in neural populations saturates as more neurons are added. Some studies have argued that it does not^35^, and that noise correlations therefore impose no fundamental bounds on coding fidelity, while others report saturation consistent with information-limiting correlations^12,36,37^. While we did not record from a large enough number of neurons to settle this issue, our results nevertheless suggests a potential mechanism that could explain the difference. We found that the alignment between the shared noise and the signal was reduced by PV-cell stimulation, and it could be that rather than a fixed property of cortical circuits, information-limiting correlations depend on behavioral states and the corresponding level of inhibition.

## METHODS

All procedures in this study conformed to the National Institutes of Health guidelines and were approved by the University of Houston Institutional Animal Care and Use Committee.

### Preparation of the animals for surgeries

Three surgeries were performed on the animals, one to implant a cranial headpost, one to implant a recording chamber, and one to inject viral vectors. In the morning of the surgery, the animals were restrained in a primate chair and sedated with an intra-muscular (IM) injection of Atropine sulfate (0.02mg/kg), followed by an IM injection of Midazolam (0.05-0.5 mg/kg) and Alfaxalone (10-12mg/kg). When the animal was sedated, it was removed from the primate chair, placed on a heating pad and covered with blankets. The animal was then intubated with a lubricated endotracheal tube, and monitoring of EKG, expired and inspired CO2, SPO2, rectal temperature and blood pressure was started. The animal’s ears and the area below the eyes were treated with a local analgesic cream (2.5% prilocaine, 2.5% lidocaine), and the animal was placed in a stereotaxic device. Isoflurane (0.1-2.5%) anesthesia was started, and the animal received an IM injection of Dexamethasone (0.5mg/kg). For IV fluids, Lactated Ringers with 5% dextrose were infused at 2-5mL/kg/hr rate. The incision site was numbed by an injection of 2% Lidocaine. The animal received a prophylactic dose of Cefazolin (22mg/kg IV) before the incision and then every 90 minutes. When necessary, isoflurane anesthesia was supplemented with Propofol (0.1-1.0 mg/kg/min). Dopamine infusion (2-10 mcg/kg/min) was used for blood pressure support during the surgery when necessary. For postoperative analgesia, Buprenorphine (0.005-0.01 mg/kg SQ/IM) and Meloxicam (0.1-0.2 mg/kg) were administered.

### Virus vectors and injections

To inject virus vectors into the visual cortex, two durotomies were performed under isoflurane anesthesia. Under surgical microscope, a glass injection pipette (tip diameter 30-50 µm) was first lowered onto the surface of the brain and then advanced to 1350 µm depth. A viral vector solution containing 180 nL of AAV.PhPeB.S5E2.ChR2-mCherry^38^ was injected into the cortex, as this vector produces up to 97% specificity to PV-cells in the marmoset visual cortex^39^.

Injections were performed at a rate of 1 nL/sec using the Nanoject III injectrode (Drummond Scientific) attached to a micromanipulator in a stereotax tower. After the injection was completed, we waited 5 minutes, after which the pipette was retracted, and the injections were repeated at 900 and 450 µm depths, at 5 minutes intervals. Upon completion of the injections, the craniotomy was filled with silastic, the chamber was closed, and the animal was recovered.

### In-vivo fluorescent imaging to monitor expression and target recordings

To target recordings to where and when the opsins are expressed, a custom epi-fluorescent imaging set-up was used. The imaging was performed with a 4x NA 0.13 objective and PCO Panda camera with 4000 ms exposure.

### Animal training

The animals were trained to target their gaze on a marmoset face, presented on a screen in front of the animal (Propixx Projector, 1920×1080 resolution, 120 Hz frame-rate, 42 cm distance from the eyes). The monkeys were rewarded for targeting their gaze on a face of a conspecific with apple juice, delivered through a tube placed above their lower lip, and controlled via a microfluidics pump. The reward volume was proportional to the duration of the fixation.

### Electrophysiology and spike-sorting

To study spiking activity in neural populations of the primate V1, recordings were performed simultaneously from 384 channels with the Neuropixels 1.0 NHP multielectrode arrays, penetrated into the cortex through the dura. The continuous voltage traces were sampled at 30kHz and saved for post-processing. To extract single and multi-unit activity, spikesorting was performed with Kilosort 4.

### Stimuli

In each session, 3-4 randomly sampled images from the Berkeley Segmentation Dataset were show windowed with a 16° diameter raised cosine and centered approximately on the receptive fields of the recorded neurons.

### Cell selection

We included units in the analyses if they satisfied three criteria. First, stimulus-evoked firing rate in control trials exceeded the pre-stimulus baseline (one-sided Wilcoxon signed-rank test, α = 0.01). Second, to exclude units that were not affected by PV-cell stimulation, laser stimulation had to significantly reduce the spike count in the 75–200 ms post-stimulus response window (one-sided Mann–Whitney U test (laser < control, α = 0.05). Third, to avoid analysing units that were directly driven by the optogenetic stimulation, we excluded cells if the laser significantly increased their firing in the 0–30 ms post-stimulus window (one-sided Mann–Whitney U, laser > control, α = 0.05). Only units passing all three criteria were carried into the analyses, and recordings with fewer than three selected units or fewer than two stimuli with ≥ 5 trials per condition were excluded.

### Computing d’ in PLS space

We used partial least squares (PLS) analysis to identify the dimensions of population spiking activity that best separates the stimuli. As optogenetic stimulation of PV-cells affects firing-rates, histogram-matching was used to control for the differences in rates (see histogram-matching). To account for the dependence between spike count variance and mean, the data was square root transformed before performing the PLS. The latent PLS space was estimated using the non-linear iterative partial least squares algorithm using the Python toolbox scikit-learn. The dimensionality of the space was selected by maximizing the binary classification accuracy with 5-fold cross-validation computed on trials. Discriminability index (d’) was computed on the low-dimensional PLS-scores that result from projecting the spike counts on the PLS axes as, 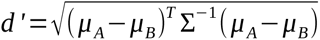, where *μ_A_* and *μ_B_* are the mean response vectors to stimulus A and B, and Σ is the pooled covariance matrix computed as trial-weighted average of the individual covariance matrices elicited by stimulus A and B. To ensure numerical stability, 10^-6^ was added to the diagonal before pseudo-inversion. Computing d′ in the low-dimensional PLS space ensures that the pooled covariance matrix can be inverted reliably even when the number of trials is comparable to or smaller than the number of neurons, a regime where the full-dimensional d’ is severely biased. Finally, we used the Kanitscheider-correction to compute bias-corrected d’ prime as,

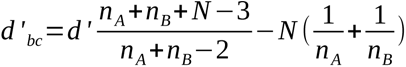, where *n_A_*and *nB* are the numbers of trials for stimulus A and B, and N is the dimensionality of the PLS space in which the d’ is computed. To understand how discriminability depends on the duration of the analysis window, d’ was computed in count windows of increasing duration.

### Covariance regularization through factor analysis

The noise covariance matrix of high-dimensional spiking activity estimated from tens of trials is under-determined and many of its entries can be dominated by sampling noise. Rather than directly using the sample covariance, we regularized it with factor analysis. For each stimulus and condition, we used the EM-algorithm to fit the model Σ=*L L^T^* +*diag* (*ψ* ), where *L*∈*ℝ^Nxd^* is a d-dimensional matrix holding the shared component of trial-to-trial spiking variability and *diag(φ)* is a diagonal matrix holding the private variability of each neuron. The latent dimensionality d was determined with 5-fold cross-validation over trials separately for the control condition and when PV-cells where stimulated. For each stimulus and condition, we used the dimensionality that maximized the log-likelihood of population spike counts in held-out trials not used for training. The Σ was normalized by 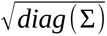 to convert covariance to correlation.

### Noise component decoders

To isolate which components of the trial-to-trial noise covariance carries the PV-cell driven change in discriminability, we built decoders that share a common signal axis but recombine noise components across conditions. The covariance matrix was decomposed using factor analysis as decribed above, as this allowed us to decompose spiking variability to private and shared components. For each image pair, factor analysis was fit separately to the within-class residuals of the control and PV stimulation conditions. For each image pair, we ran 50 bootstrap iterations to resample trials and the histogram matched neuron subsets. The FA dimensionality was selected within every bootstrap by cross validation and FA was then fit with the selected dimensionality on 5 train/test splits. Discriminability was evaluated against control anchored signal axis ŝ = Δµ_control, train_/‖Δµ_control, train_‖, computed strcitly from the training split. d-prime was computed as 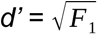 where F1 is the 1D linear Fisher information computed as 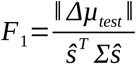 , and Σ was constructed from different combinations of private and shared variability. To dissociate PV-mediated reorientation of the share noise subspace from change in the noise spectrum, we used two further decompositions of Σ, one in which the eigenvectors were estimated from the stimulation trials and the spectrum from the control trials, and vice versa.

## ACKNOWLEDGMENTS

We are grateful to Dr. Jude Mitchell and Dr. Nicholas Priebe for providing animals for the project, and for Ms. Colleen LaBorde for excellent administrative support. The project was funded by the following grants awarded to LN, NIH R00 EY029374, Kavli Foundation Seed Grant, University of Houston Research Excellence Award and start-up funds from the University of Houston Division of Research and College of Optometry.

## REFERENCES

1. Fechner, G. Elemente Der Psychophysik. vol. 2 (Breitkopf und Härtel, Leipzig, 1860).

2. Green, D. & Swets, J. Signal Detection Theory. (Wiley, New York, 1966).

3. Goris, R. L. T., Movshon, J. A. & Simoncelli, E. P. Partitioning neuronal variability. Nat. Neurosci. 17, 858–865 (2014).

4. Tolhurst, D. J., Movshon, J. A. & Dean, A. F. The statistical reliability of signals in single neurons in cat and monkey visual cortex. Vision Res. 23, 775–785 (1983).

5. Zohary, E., Shadlen, M. N. & Newsome, W. T. Correlated neuronal discharge rate and its implications for psychophysical performance. Nature 370, 140–143 (1994).

6. Shadlen, M., Britten, K., Newsome, W. & Movshon, J. A computational analysis of the relationship between neuronal and behavioral responses to visual motion. J. Neurosci. 16, 1486–1510 (1996).

7. Shadlen, M. N. & Newsome, W. T. The Variable Discharge of Cortical Neurons: Implications for Connectivity, Computation, and Information Coding. J. Neurosci. 18, 3870–3896 (1998).

8. Gutnisky, D. A. & Dragoi, V. Adaptive coding of visual information in neural populations. Nature 452, 220–224 (2008).

9. Wakhloo, A. J., Slatton, W. & Chung, S. Neural population geometry and optimal coding of tasks with shared latent structure. Nat. Neurosci. 29, 682–692 (2026).

10. Averbeck, B. B., Latham, P. E. & Pouget, A. Neural correlations, population coding and computation. Nat. Rev. Neurosci. 7, 358–366 (2006).

11. Moreno-Bote, R. et al. Information-limiting correlations. Nat. Neurosci. 17, 1410–1417 (2014).

12. Rumyantsev, O. I. et al. Fundamental bounds on the fidelity of sensory cortical coding. Nature 580, 100–105 (2020).

13. Mitchell, J. F., Sundberg, K. A. & Reynolds, J. H. Differential Attention-Dependent Response Modulation across Cell Classes in Macaque Visual Area V4. Neuron 55, 131–141 (2007).

14. Mitchell, J. F., Sundberg, K. A. & Reynolds, J. H. Spatial Attention Decorrelates Intrinsic Activity Fluctuations in Macaque Area V4. Neuron 63, 879–888 (2009).

15. Cohen, M. R. & Maunsell, J. H. R. Attention improves performance primarily by reducing interneuronal correlations. Nat. Neurosci. 12, 1594–1600 (2009).

16. Merrikhi, Y., Clark, K. & Noudoost, B. Concurrent influence of top-down and bottom-up inputs on correlated activity of Macaque extrastriate neurons. Nat. Commun. 9, 5393 (2018).

17. Ni, A. M., Ruff, D. A., Alberts, J. J., Symmonds, J. & Cohen, M. R. Learning and attention reveal a general relationship between population activity and behavior. Science 359, 463–465 (2018).

18. Hennequin, G., Ahmadian, Y., Rubin, D. B., Lengyel, M. & Miller, K. D. The Dynamical Regime of Sensory Cortex: Stable Dynamics around a Single Stimulus-Tuned Attractor Account for Patterns of Noise Variability. Neuron 98, 846–860.e5 (2018).

19. Renart, A. et al. The Asynchronous State in Cortical Circuits. Science 327, 587–590 (2010).

20. Rosenbaum, R., Smith, M. A., Kohn, A., Rubin, J. E. & Doiron, B. The spatial structure of correlated neuronal variability. Nat. Neurosci. 20, 107–114 (2017).

21. Huang, C. et al. Circuit Models of Low-Dimensional Shared Variability in Cortical Networks. Neuron 101, 337–348.e4 (2019).

22. Kanitscheider, I., Coen-Cagli, R., Kohn, A. & Pouget, A. Measuring Fisher Information Accurately in Correlated Neural Populations. PLOS Comput. Biol. 11, e1004218 (2015).

23. Smith, M. A. & Kohn, A. Spatial and Temporal Scales of Neuronal Correlation in Primary Visual Cortex. J. Neurosci. 28, 12591–12603 (2008).

24. Yu, B. M. et al. Gaussian-process factor analysis for low-dimensional single-trial analysis of neural population activity. in Advances in Neural Information Processing Systems (eds Koller, D., Schuurmans, D., Bengio, Y. & Bottou, L.) vol. 21 (Curran Associates, Inc., 2008).

25. Semedo, J. D., Zandvakili, A., Machens, C. K., Yu, B. M. & Kohn, A. Cortical Areas Interact through a Communication Subspace. Neuron 102, 249–259.e4 (2019).

26. Nurminen, L., Bijanzadeh, M. & Angelucci, A. Size tuning of neural response variability in laminar circuits of macaque primary visual cortex. eLife 15, e86334 (2026).

27. Snyder, A. C., Morais, M. J., Kohn, A. & Smith, M. A. Correlations in V1 Are Reduced by Stimulation Outside the Receptive Field. J. Neurosci. 34, 11222–11227 (2014).

28. Festa, D., Aschner, A., Davila, A., Kohn, A. & Coen-Cagli, R. Neuronal variability reflects probabilistic inference tuned to natural image statistics. Nat. Commun. 12, 3635 (2021).

29. Stringer, C. et al. Inhibitory control of correlated intrinsic variability in cortical networks. eLife 5, e19695 (2016).

30. Atallah, B. V., Bruns, W., Carandini, M. & Scanziani, M. Parvalbumin-Expressing Interneurons Linearly Transform Cortical Responses to Visual Stimuli. Neuron 73, 159–170 (2012).

31. Vafaei, A. et al. Laminar-specific control of response gain and orientation-tuning by parvalbumin-expressing inhibitory interneurons in primate visual cortex. Preprint at 10.64898/2025.12.23.696300 (2025).

32. Coen-Cagli, R. & Solomon, S. S. Relating Divisive Normalization to Neuronal Response Variability. J. Neurosci. 39, 7344–7356 (2019).

33. Hofer, S. B. et al. Differential connectivity and response dynamics of excitatory and inhibitory neurons in visual cortex. Nat. Neurosci. 14, 1045–1052 (2011).

34. Znamenskiy, P. et al. Functional specificity of recurrent inhibition in visual cortex. Neuron 112, 991–1000.e8 (2024).

35. Stringer, C., Michaelos, M., Tsyboulski, D., Lindo, S. E. & Pachitariu, M. High-precision coding in visual cortex. Cell 184, 2767–2778.e15 (2021).

36. Bartolo, R., Saunders, R. C., Mitz, A. R. & Averbeck, B. B. Information-Limiting Correlations in Large Neural Populations. J. Neurosci. 40, 1668–1678 (2020).

37. Kafashan, M. et al. Scaling of sensory information in large neural populations shows signatures of information-limiting correlations. Nat. Commun. 12, 473 (2021).

38. Vormstein-Schneider, D. et al. Viral manipulation of functionally distinct interneurons in mice, non-human primates and humans. Nat. Neurosci. 23, 1629–1636 (2020).

39. Federer, F. et al. Laminar specificity and coverage of viral-mediated gene expression restricted to GABAergic interneurons and their parvalbumin subclass in marmoset primary visual cortex. eLife 13, RP97673 (2024).

40. Livezey, J. A., Sachdeva, P. S., Dougherty, M. E., Summers, M. T. & Bouchard, K. E. The geometry of correlated variability leads to highly suboptimal discriminative sensory coding. J. Neurophysiol. 133, 124–141 (2025).

